# Repeated evolution of bedaquiline resistance in *Mycobacterium tuberculosis* is driven by truncation of *mmpR5*

**DOI:** 10.1101/2022.12.08.519610

**Authors:** Leah W Roberts, Kerri M Malone, Martin Hunt, Lavania Joseph, Penelope Wintringer, Jeff Knaggs, Derrick Crook, Maha R Farhat, Zamin Iqbal, Shaheed V Omar

## Abstract

The antibiotic Bedaquiline (BDQ) is a key component of new WHO regimens for drug resistant tuberculosis (TB) but predicting BDQ resistance (BDQ-R) from genotypes remains challenging. We analysed a collection (n=505) of *Mycobacterium tuberculosis* from two high prevalence areas in South Africa (Cape Town and Johannesburg, 2019-2020), and found 53 independent acquisitions of 31 different mutations within the *mmpR5* regulatory gene, with a particular enrichment of truncated MmpR5 in BDQ-R isolates by either frameshift or introduction of an insertion element. Truncations occurred across three *M. tuberculosis* lineages, impacting 66% of BDQ-R isolates. Extending our analysis to 1,961 isolates with minimum inhibitory concentrations (MICs) revealed that *mmpR5*-disrupted isolates had a median BDQ MIC of 0.25 mg/L, compared to the wild-type median of 0.06 mg/L. By matching *mmpR5*-disrupted isolates with phylogenetically close control isolates without the disruption, we were able to estimate the impact on MIC of individual mutations. In conclusion, as the MIC increase borders the ECOFF threshold for BDQ-R, we recommend the continued use of MICs and detection of MmpR5 truncations to identify modest shifts in BDQ-R.

## Introduction

Tuberculosis (TB), caused by *Mycobacterium tuberculosis*, is one of the oldest transmissible diseases in human history and remains a public health priority. While the incidence of TB in developed countries has improved substantially (to less than 1 death per 100,000 population per year), the global burden remains high, with an estimated 10 million infections and 1.5 million deaths annually (WHO 2022 Report). Our ability to treat TB is limited by the ability of *M. tuberculosis* to develop resistance to prescribed treatment regimens over time. As such, rapid understanding of the *M. tuberculosis* resistance profile is critical to administering the best drug combination for successful treatment.

Bedaquiline (BDQ) is a new TB drug that was approved for use in 2012; the first new TB drug in more than 40 years^1^. BDQ is an ATP synthase inhibitor with strong bacteriocidal activity that has been successfully used in combination to treat rifampicin-resistant/multidrug-resistant TB (RR/MDR-TB)^2,3^. Early BDQ usage was conservative in cases where other treatment strategies were ineffective or unavailable. However, recent evidence has led to the World Health Organisation (WHO) recommending BDQ as part of combination therapies (such as the BPaLM regimen) to treat RR/MDR-TB (WHO May 2022). As of 2022, 124 countries were using BDQ as part of treatment for drug-resistant TB (DR-TB) (WHO Report 2022).

In 2020, eight countries contributed two thirds of the global TB incidence (WHO report 2021). One of these countries, South Africa, accounted for 3.3% of the global total, and was one of 30 countries listed by the WHO to have high rates of RR/MDR-TB (WHO 2021 report). Routine use of BDQ began in South Africa in 2015 and accounts for over >50% of global BDQ use by 2020^4^. Although BDQ has been widely successful in reducing mortality in patients with RR/MDR-TB^5,6^, emerging resistance threatens to hamper the successful treatment and management of TB globally.

Several studies have attempted to identify the genetic determinants of BDQ resistance. One of the earliest reports of resistance identified a target-based mutation in the gene *atpE*, however, mutations in *atpE* are rarely seen *in vivo* due to deleterious effects to overall fitness^2^. Further studies have since identified the regulatory gene *mmpL5* (or *rv0678*) as a common site for mutations in BDQ-R isolates^4,7,8^. The translated protein MmpR5 is a 165 amino acid (aa) transcriptional repressor of the efflux pump MmpS5-MmpL5, which has been associated with resistance to both BDQ and clofazimine. So far, no single mutation in *mmpR5* appears strongly linked to BDQ resistance in *M. tuberculosis* isolates^9–11^, with most leading to only modest MIC increases between 2- and 4-fold^10^. Mutations in *mmpR5* have also been found without prior BDQ or clofazimine treatment^12^, suggesting that they are naturally occurring.

Here, we use whole genome sequencing (WGS) to analyse a collection of *M. tuberculosis* sampled randomly from patients in Johannesburg and Cape Town (South Africa) between 2019-2020. We use local assembly to show that truncation of *mmpR5* via several mechanisms explains almost 70% of BDQ-R isolates. We further extend our study to include all publicly available WGS of South African *M. tuberculosis* to date, including a set of ∼2,000 phenotyped *M. tuberculosis* from the CRyPTIC consortium^13^. Assessment of isolate MICs in relation to MmpR5 truncation showed a clear shift in BDQ susceptibility, while truncation of MmpL5 produced a hypersensitive phenotype. As this shift in BDQ MIC is unrecognisable with binary phenotypes, a combination of MICs and MmpR5 truncations could give more insight into BDQ resistance in *M. tuberculosis*.

## Results

We collected 505 *M. tuberculosis* samples from 483 patients in South Africa between 2019 to 2020 (hereafter referred to as **SA1**). *In silico* antimicrobial resistance (AMR) typing found the majority of isolates to be multidrug-resistant (MDR) with resistance to both rifampicin and isoniazid but susceptible to fluoroquinolones (n=347, 69%) (**Figure 1**). A small number of isolates were rifampicin-mono resistance (RR-TB; n=17, 3%) or isoniazid-mono resistant (IR-TB; n=7, 1%). 126 isolates (25%) were pre-extensively drug-resistant (pre-XDR) with MDR-profiles plus resistance to fluoroquinolones. Seven isolates were susceptible to rifampicin, isoniazid, and fluoroquinolone drugs. A single isolate was isoniazid- and fluoroquinolone-resistant (including moxifloxacin), but susceptible to rifampicin. Comparison of *in silico* typing to available phenotypes showed 84-96% concordance between phenotypic and genotypic predictions (**supplementary dataset 1**).

**Figure 1:**
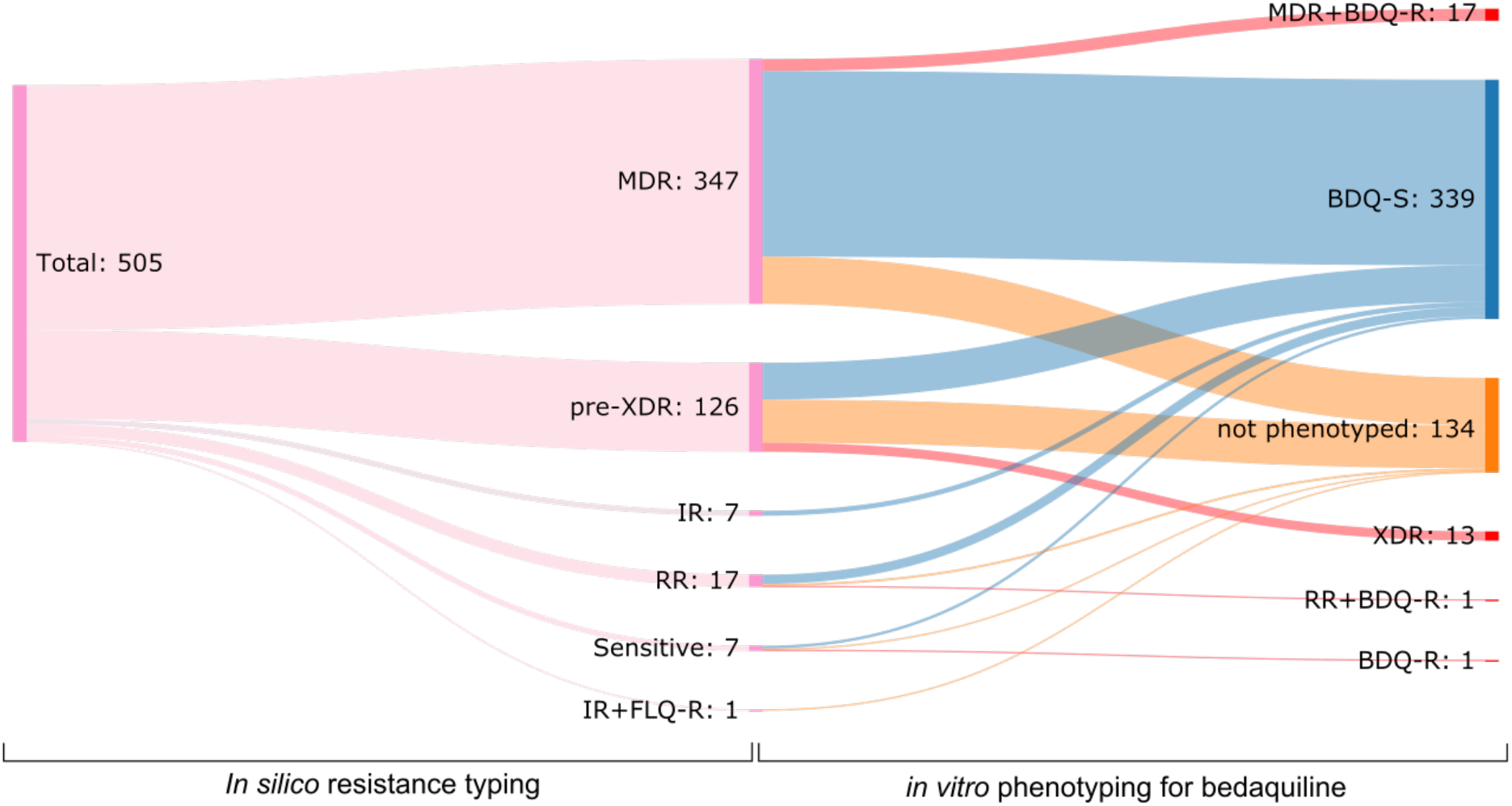
Antimicrobial resistance typing using *in silico* and phenotypic methods. AMR profiles were determined for each *M. tuberculosis* isolate in the SA1 dataset using Mykrobe (left panel). A subset were also phenotyped for resistance to bedaquiline (BDQ) (right panel). The right panel adds phenotypic information (*i*.*e*. bedaquiline susceptibility) to the *in silico* resistance typings (*e*.*g*. a pre-XDR isolate with phenotypic bedaquiline resistance would then be classed as XDR). Abbreviations are consistent with the WHO guidelines for *M. tuberculosis* resistance typing. IR = isoniazid resistant, RR = rifampicin resistant, BDQ-R = bedaquiline resistant, XDR = extensively drug resistant, MDR = multidrug resistant.

A random subset of the SA1 dataset (n=371) was available at the time for phenotyping, which identified 32 isolates resistant to bedaquiline (BDQ-R). When combined with the *in silico* AMR predictions, 13 (10%) pre-XDR isolates could be reclassified as extensively-drug resistant (XDR), while 52 remained pre-XDR (41%), and 61 were not tested (49%) (**Figure 1**). Additionally, 17 MDR-TB, one RR-TB and one sensitive TB also had BDQ-R phenotypes.

To predict how BDQ resistance emerged (clonally from a single event, or multiple times independently), we first created a maximum-likelihood phylogeny using core single nucleotide polymorphisms (SNPs) from the SA1 dataset. This revealed a mix of lineages, with lineage 2 the most common (n=314), followed by lineage 4 (n=178), lineage 1 (n=10) and lineage 3 (n=3). Very few incidents of recent transmission (defined as within 5 SNPs) were identified (13 incidents involving 33 samples) (**Figure 2**), although there were notable clonal expansions within lineages 2.2 and 2.2.1 (**detailed further in the supplementary materials**). Phenotypically BDQ-R isolates were spread throughout the phylogeny, where we predict at least 28 independent events leading to resistance (**Figure 2**).

**Figure 2:**
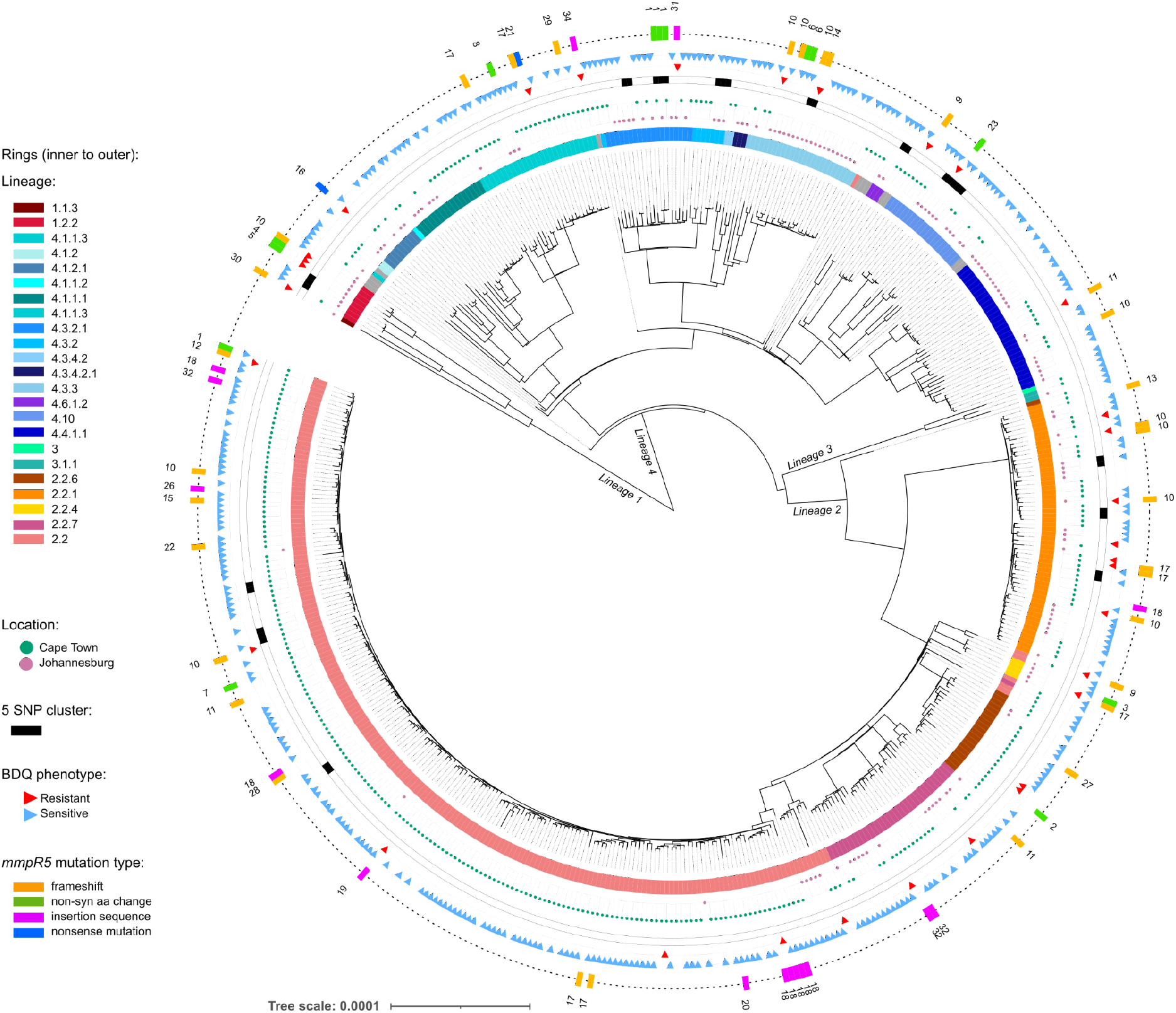
SA1 dataset from Cape Town and Johannesburg collected between 2019-2020. Phylogeny built using core SNPs with IQTree v1.6.12. Outer ring (with numbers) denotes *mmpR5* variant identifier as used in **Table 1**.

### BDQ-R is associated with truncation of the regulatory gene *mmpR5*

To identify mutations that could be associated with BDQ-R, we investigated two genes previously reported to have a role in BDQ-R: the target gene *atpE* and the efflux pump regulator *mmpR5*. None of the isolates in SA1 had variants in *atpE*. Additionally, the vast majority of isolates had no variants in *mmpR5* (n=441, 87%). However, 13 isolates (3%) had non-synonymous mutations in *mmpR5*, of which three were BDQ-R isolates (G41D, n=1; R72W, n=1; L117V n=1; note 4/13 were not phenotyped) (**Table 1**). Of the remainder (n=51), 33 were found to have frameshift mutations. In order to interpret the effect of these frameshifts, we obtained the predicted protein sequence for *mmpR5* for all isolates (using *de novo* assemblies) to assess possible truncation events. Overall, we identified 48 isolates (9.5%) with a truncated MmpR5 protein (**Figure 3**).

**Table 1:** Summary of variants identified in *mmpR5* (SA1 dataset)

| mutation number | isolate | accession | type | mutation | BDQ phenotype | aa length |
| --- | --- | --- | --- | --- | --- | --- |
| 1 | S006 | ERS14298562 | non-syn aa | M139I | n.a. | 165 |
| 1 | S025 | ERS14298581 | non-syn aa | M139I | n.a. | 165 |
| 1 | S040 | ERS14298596 | non-syn aa | M139I | n.a. | 165 |
| 1 | S435 | ERS14298991 | non-syn aa | M139I | S | 165 |
| 2 | S063 | ERS14298619 | non-syn aa | A99V | n.a. | 165 |
| 3 | S135 | ERS14298691 | non-syn aa | R82W | S | 165 |
| 4 | S145 | ERS14298701 | non-syn aa | G41D | R | 165 |
| 5 | S198 | ERS14298754 | non-syn aa | R72W | R | 165 |
| 6 | S199 | ERS14298755 | non-syn aa | F100I | S | 165 |
| 6 | S200 | ERS14298756 | non-syn aa | F100I | S | 165 |
| 7 | S392 | ERS14298948 | non-syn aa | L95S | S | 165 |
| 8 | S454 | ERS14299010 | non-syn aa | F19V | S | 165 |
| 9 | S165 | ERS14298721 | frameshift | V45fs | R | 45 |
| 9 | S387 | ERS14298943 | frameshift | V45fs | R | 45 |
| 10 | S044 | ERS14298600 | frameshift | P48fs:RAAVL | n.a. | 80 |
| 10 | S094 | ERS14298650 | frameshift | P48fs:RAAVL | n.a. | 80 |
| 10 | S125 | ERS14298681 | frameshift | P48fs:RAAVL | R | 80 |
| 10 | S134 | ERS14298690 | frameshift | P48fs:RAAVL | R | 80 |
| 10 | S172 | ERS14298728 | frameshift | P48fs:RAAVL | n.a. | 80 |
| 10 | S194 | ERS14298750 | frameshift | P48fs:RAAVL | n.a. | 80 |
| 10 | S257 | ERS14298813 | frameshift | P48fs:RAAVL | S | 80 |
| 10 | S279 | ERS14298835 | frameshift | P48fs:RAAVL | R | 80 |
| 10 | S393 | ERS14298949 | frameshift | P48fs:RAAVL | R | 80 |
| 10 | S425 | ERS14298981 | frameshift | P48fs:RAAVL | R | 80 |
| 10 | S426 | ERS14298982 | frameshift | P48fs:RAAVL | R | 80 |
| 11 | S033 | ERS14298589 | frameshift | G66fs:DQHQC | n.a. | 80 |
| 11 | S059 | ERS14298615 | frameshift | G66fs:DQHQC | R | 80 |
| 11 | S067 | ERS14298623 | frameshift | G66fs:DQHQC | n.a. | 80 |
| 11 | S082 | ERS14298638 | frameshift | G66fs:DQHQC | n.a. | 80 |
| 12 | S255 | ERS14298811 | frameshift | C46fs:GSRAA | R | 80 |
| 13 | S068 | ERS14298624 | frameshift | C46fs:DPERP | n.a. | 80 |
| 14 | S176 | ERS14298732 | frameshift | C46fs:VIPSG | R | 74 |
| 15 | S232 | ERS14298788 | frameshift | G66fs:RSAPM | S | 74 |
| 16 | S177 | ERS14298733 | nonsense | W42nonsense | R | 41 |
| 17 | S056 | ERS14298612 | frameshift | G66fs:SAPMP | n.a. | 73 |
| 17 | S085 | ERS14298641 | frameshift | G66fs:SAPMP | n.a. | 73 |
| 17 | S141 | ERS14298697 | frameshift | G66fs:SAPMP | R | 73 |
| 17 | S183 | ERS14298739 | frameshift | G66fs:SAPMP | R | 73 |
| 17 | S184 | ERS14298740 | frameshift | G66fs:SAPMP | S | 73 |
| 17 | S203 | ERS14298759 | frameshift | G66fs:SAPMP | n.a. | 73 |
| 17 | S300 | ERS14298856 | frameshift | G66fs:SAPMP | S | 73 |
| 18 | S011 | ERS14298567 | IS insertion | IS6110:779036 | n.a. | 16 |
| 18 | S088 | ERS14298644 | IS insertion | IS6110:779043 | n.a. | 21 |
| 18 | S124 | ERS14298680 | IS insertion | IS6110:779036 | S | 21 |
| 18 | S295 | ERS14298851 | IS insertion | IS6110:779041 | S | 21 |
| 18 | S370 | ERS14298926 | IS insertion | IS6110:779046 | n.a. | 21 |
| 18 | S376 | ERS14298932 | IS insertion | IS6110:779045 | R | 21 |
| 18 | S499 | ERS14299055 | IS insertion | IS6110:779045 | S | 21 |
| 18 | S020 | ERS14298576 | IS insertion | IS6110:779042 | n.a. | 33 |
| 19 | S321 | ERS14298877 | IS insertion | IS6110:779075 | R | 29 |
| 20 | S432 | ERS14298988 | IS insertion | IS6110:779056 | S | 25 |
| 21 | S449 | ERS14299005 | nonsense | Q76nonsense | R | 75 |
| 22 | S456 | ERS14299012 | frameshift | G65fs | S | 65 |
| 23 | S472 | ERS14299028 | Non-syn aa +<br>unknown truncation* | L117V:R135stop | R | 134 |
| 26 | S042 | ERS14298598 | IS insertion | IS6110:779026 | n.a. | 11 |
| 27 | S057 | ERS14298613 | frameshift | D47fs:LPSGS | n.a. | 74 |
| 28 | S075 | ERS14298631 | frameshift | P48fs | n.a. | 40 |
| 29 | S092 | ERS14298648 | frameshift | R89fs | n.a. | 105 |
| 30 | S095 | ERS14298651 | frameshift | G6fs | R | 31 |
| 31 | S108 | ERS14298664 | IS insertion | IS6110:779168 | R | 162 |
| 32 | S043 | ERS14298599 | IS promoter | IS6110:778978 | n.a. | 165 |
| 32 | S215 | ERS14298771 | IS promoter | IS6110:778978 | R | 165 |
| 32 | S254 | ERS14298810 | IS promoter | IS6110:778978 | S | 165 |
| 34 | S412 | ERS14298968 | IS promoter | IS6110:778945 | R | 165 |

\* This isolate was not included in results relating to MmpR5 truncation

**Figure 3:**
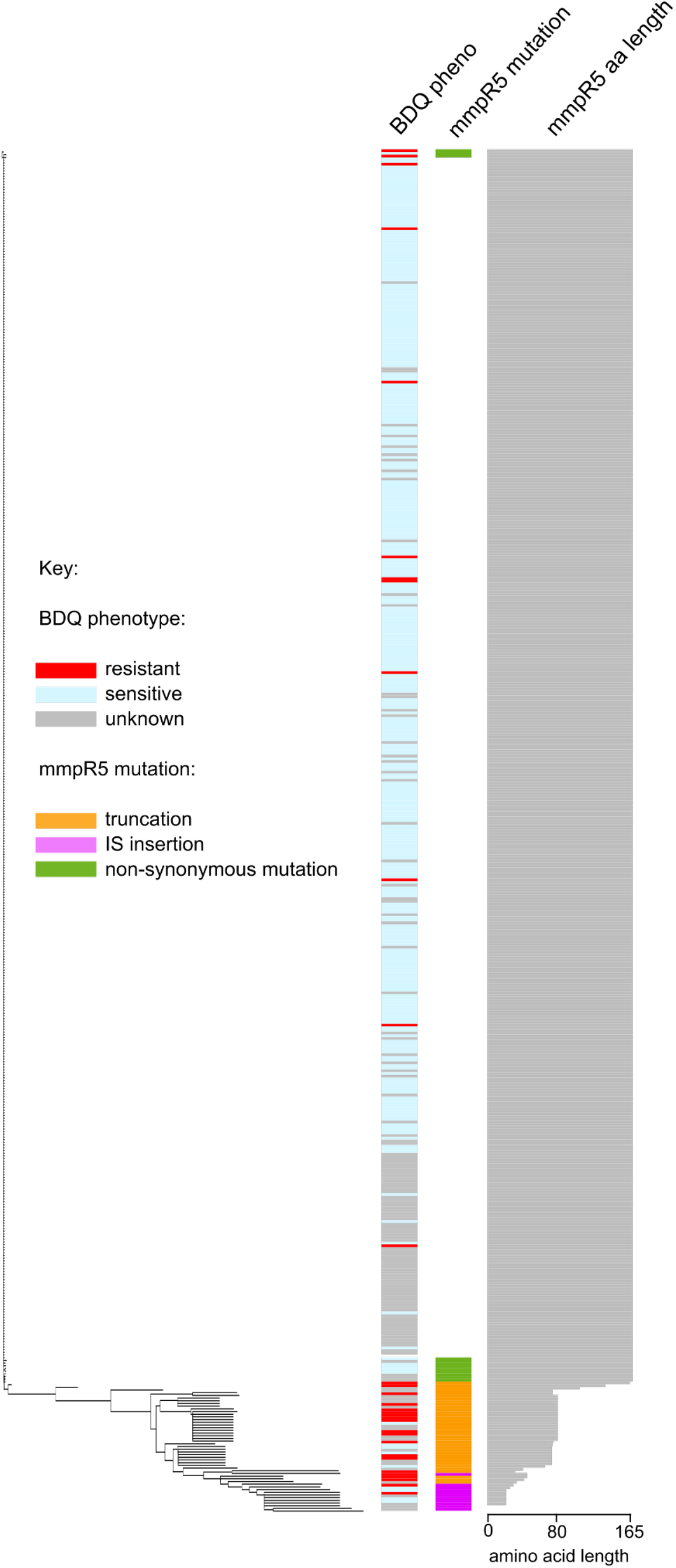
protein alignment and neighbour joining tree of MmpR5 predicted protein sequences from the SA1 dataset

The length of truncation varied from 3 to 154 amino acids (from a total length of 165 amino acids for MmpR5). 33/48 isolates were found to have one of 14 different frameshift mutations (**Table 1**). Further inspection of the *de novo* assemblies revealed a number of isolates with suspected insertion sequences (IS) disruptions. To investigate this, we ran ISmapper^14^ looking for IS6110 insertions within or upstream of *mmpR5*. We found 12 isolates with IS6110 interrupting *mmpR5* (at 5 sites), and 4 with IS6110 in the promoter region (at 2 sites). Two isolates had different nonsense mutations, and one isolate had an unknown truncation (which we did not include further; see Methods) (**Table 1**).

Based on isolates with BDQ phenotypes (n=371), mutation and truncation events in *mmpR5* were more common in BDQ-R isolates (**Figure 3**). 24/32 (75%) resistant isolates had mutations in *mmpR5*, of which 19 (79%) were predicted to result in MmpR5 truncations. Only 5% (16/339) of sensitive isolates had mutations in *mmpR5*, of which just 9 resulted in predicted MmpR5 truncations. There was little difference in the types of mutations leading to truncations between BDQ-R and BDQ-S isolates. Frameshifts were responsible for the most MmpR5 truncations (5 in BDQ-S isolates, 14 in BDQ-R isolates). IS6110 was responsible for 4 truncations in BDQ-S isolates and 3 in BDQ-R isolates. The remainder of truncations were caused by nonsense mutations (n=2).

Comparison of each unique *mmpR5* mutation (**Table 1**) to the phylogeny revealed mostly independent acquisitions of the same mutations relatively recently in the tree (**Figure 2**). Of the 64 isolates with *mmpR5* mutations (including promoter region IS), we estimate 53 independent acquisitions (53/64=83%) of 31 different *mmpR5* mutations.

Overall, 21/32 (66%) BDQ-R isolates in our dataset had evidence of truncation (*i*.*e*. shortened proteins) or promoter interruptions in *mmpR5*. However, despite being able to correlate a large number of BDQ-R isolates with disruptions in *mmpR5*, we still observed cases where *mmpR5* was truncated but the BDQ phenotype remained sensitive, or BDQ-R isolates with an unaltered *mmpR5* gene. We therefore investigated a number of other genes that have previously been implicated in BDQ-R mechanisms in *M. tuberculosis* (*mmpL5, mmpL3, mmpS5, amiA2, pepQ, era, rv1816, rv3249c*). We found a few lineage-specific mutations (**supplementary figure 1**), but no mutations that further explained the BDQ-R isolates. To check for gene duplications of *mmpR5, mmpL5* and *mmpS5*, we also mapped all reads to the reference genome H37Rv and compared the average genome coverage to coverage at these specific genes. We found no change in coverage that could suggest duplication of these genes in any isolates. Finally, we looked for IS interruption of *mmpL5, mmpS5* or *atpE*, and found no evidence of IS6110 insertion.

### Truncation of mmpR5 was observed in all lineages across a large South African dataset

To contextualise our SA1 dataset into a broader South African cohort with accompanying MICs, we downloaded all publicly available South African *M. tuberculosis* isolates from the ENA and CRyPTIC consortium^13^ datasets (n=5,253; hereafter referred to as “SA2”). Overall, we found that our SA1 dataset was well-sampled across the known diversity of *M. tuberculosis* in South Africa, with the exception of a large lineage 2.2 clade primarily isolated in Cape Town (as noted previously; **Figure 4**).

**Figure 4.**
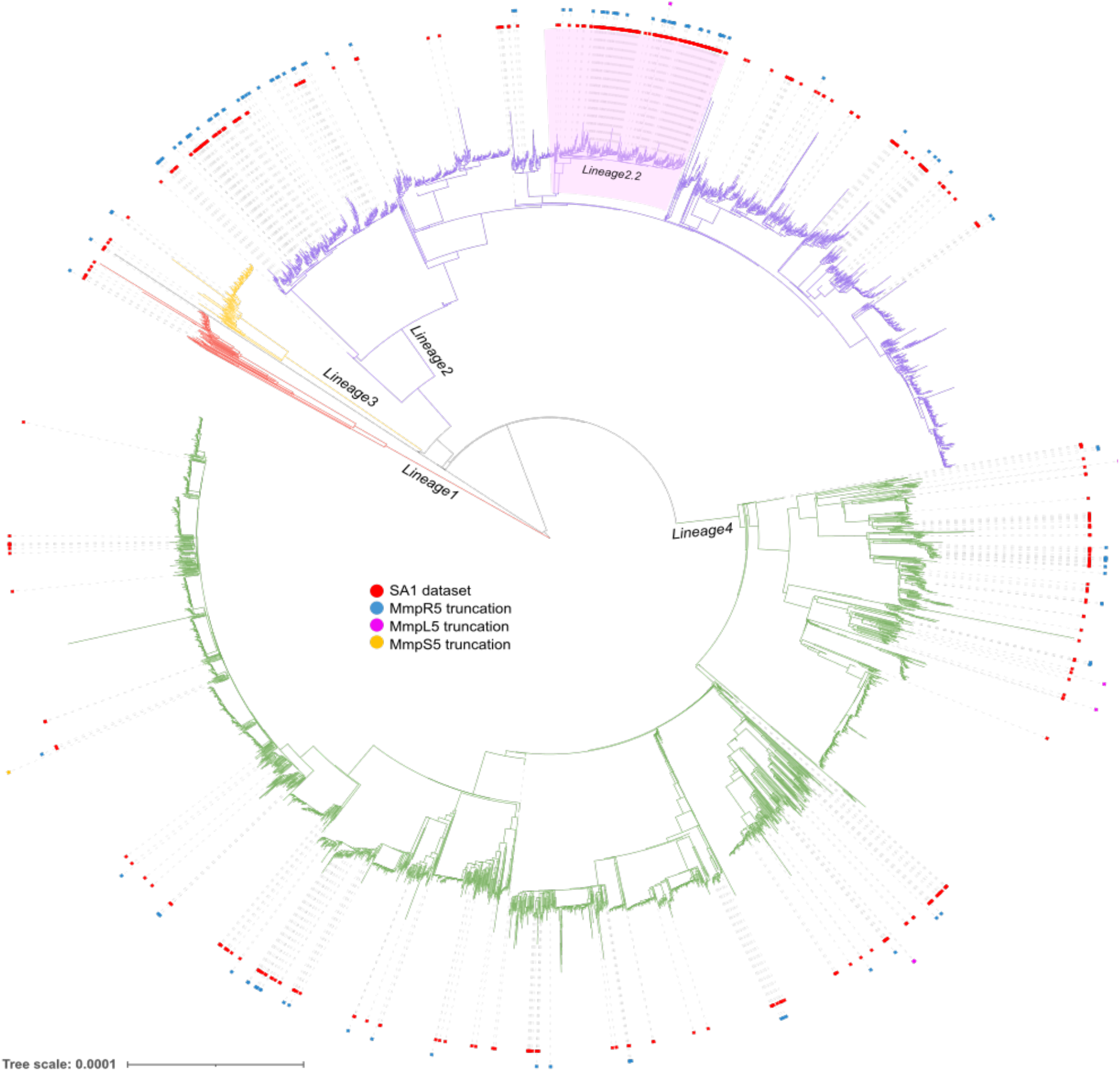
Maximum likelihood phylogeny of 5,253 publicly available South African *M. tuberculosis* isolates (SA2) plus the SA1 dataset, built using core SNPs with IQTree v2.2.0.

We used the antimicrobial resistance prediction tool ARIBA^15^ with a custom reference gene set to rapidly identify truncated genes, including *mmpR5*. ARIBA was able to identify 42/47 gene truncations (89%) from our exhaustively manually curated SA1 dataset. Three of the five missing calls were caused by IS insertions, and two had ambiguous heterogeneous calls that ARIBA did not report. Nevertheless, as ARIBA captured the majority of truncations, we used it to identify non-synonymous mutations and truncations in *mmpR5, mmpL5* and *mmpS5* in the much larger SA2 dataset.

We identified 76 isolates in the SA2 dataset with predicted truncations resulting from 18 mutations, 11 of which were unique to the SA2 dataset (including 7 frameshifts, 1 IS6110 insertion, and 3 nonsense mutations) (**supplementary dataset 1**). Three additional isolates had one of two different frameshift mutations that resulted in altered proteins but not truncation. Plotting truncations onto the large SA1+SA2 phylogeny produced the same scattered prevalence as with the smaller SA1 dataset, confirming that truncations of MmpR5 occurred independently (**Figure 4**).

We also identified 50 isolates with one of 30 additional non-synonymous variant combinations in MmpR5. When assessing all variants in *mmpR5* across both datasets, 43 mutations were unique to the SA2 dataset, eight mutations were found in both the SA1 and SA2 datasets, while 20 were unique to the SA1 dataset (not including the IS promoter mutations, which were not identified in the SA2 dataset).

Across the SA1 and SA2 datasets we identified 73 separate variants in *mmpR5* (**supplementary dataset 1**). 48/73 variants were represented by only a single isolate, while 9/73 were represented by only 2 isolates (**Figure 5; B**). 15 variant sites represented the majority of mutations, capturing 125/194 isolates with mutations in *mmpR5* (**Figure 5; A**). The most common mutations were all frameshifts at positions C46, G66 and P48, representing 40% (n=77/194) of the isolates (**Figure 5; A**). There was also a concentration of amino acid substitutions at the final “turn” of the predicted MmpR5 protein. In relation to the functional domains of the protein, we observed that the more common mutations occur in the first half of the predicted protein, and resulted in removal or disruption of the DNA-binding domain (**Figure 5; C, Supplementary Figure 2**).

**Figure 5:**
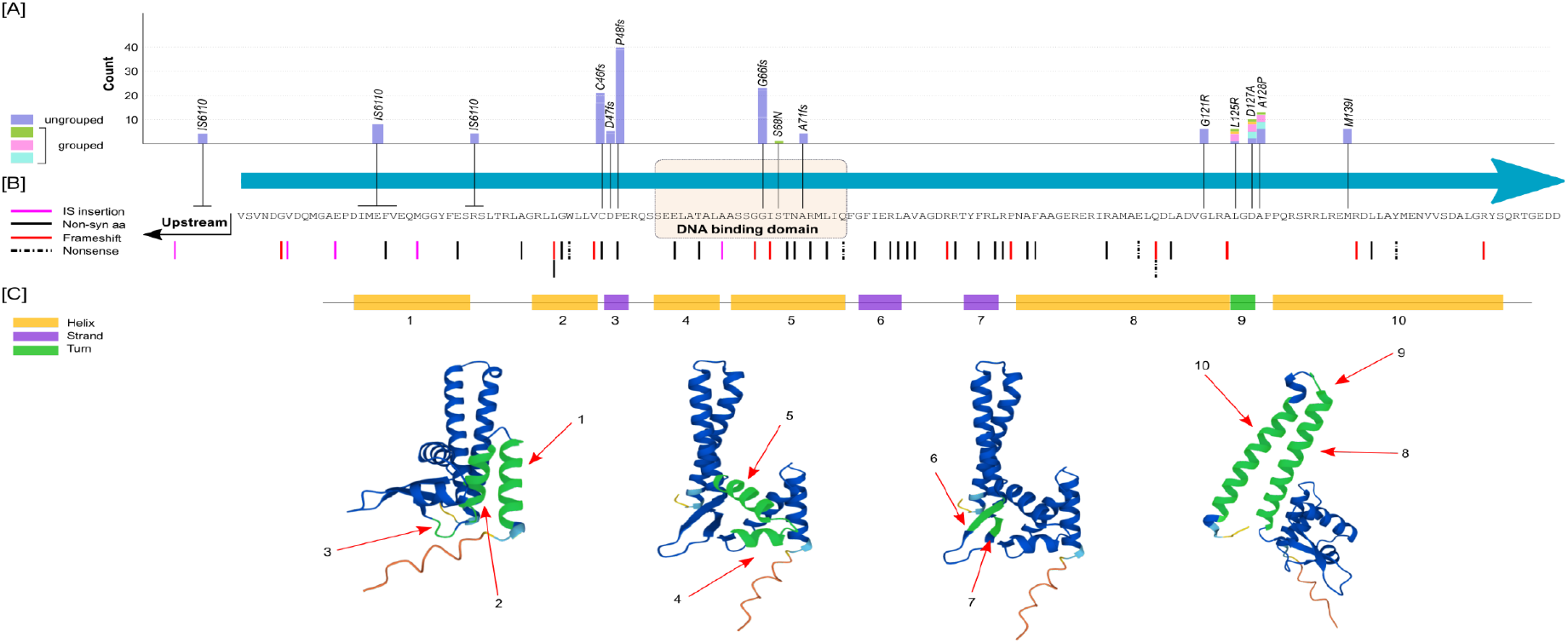
Summary of *mmpR5* mutations: [A]: all mutations that occurred in >=3 isolates. Most mutations were found in separate samples (purple bars; ungrouped). Green, light blue and pink bars represent mutations found together in a single sample (grouped). [B]: all other mutations found in <=2 isolates, and [C]: Uniprot/AlphaFold prediction of protein structure (https://www.uniprot.org/uniprotkb/I6Y8F7/entry accessed 19-08-2022)

In addition to *mmpR5* truncations, we also looked for evidence of truncation in *mmpL5* and *mmpS5* in the SA2 dataset. Overall we found 6 isolates with MmpL5 truncations, caused by 5 different mutations (Q333fs, Q702nonsense, R734fs, R115fs, and G138fs). There was no evidence of IS6110 interruption within or upstream of *mmpL5*. We only identified a single isolate with a mutation leading to a truncation in *mmpS5* (P101fs). Importantly, we did not find any isolates that had multiple gene truncations involving any combination of *mmpL5, mmpR5* or *mmpS5*. Five of the six mmpL5 truncations were phylogenetically distinct and the result of independent acquisitions (**Figure 4**). We discuss the phenotypic impact of these below.

### The BDQ minimum inhibitory concentration (MIC) is increased in isolates with mmpR5 truncation

Of the 79 MmpR5-truncated (or otherwise frameshifted) isolates found in the SA2 dataset, 75 were from the CRyPTIC dataset and had phenotypic data. 51 (68%) were BDQ-R (based on an ECOFF threshold of 0.25 mg/L for resistance), while the remainder were BDQ-S. Four of the six isolates with MmpL5 truncations had phenotypic data and were all sensitive. The single isolate with MmpS5 truncated had no phenotypic data.

As before, we found a number of BDQ-S isolates with predicted MmpR5 truncations. To gain better insight into the effect of MmpR5 truncation on resistance beyond the binary phenotypes, we plotted the BDQ minimum inhibitory concentrations (MIC) for 1,961 South African CRyPTIC isolates and separated them based on predicted truncation of MmpR5 and MmpL5 (no MIC was available for the MmpS5 truncated isolate). We found that the median BDQ MIC was elevated in isolates with MmpR5 truncated (0.25 mg/L, [stdev 0.22 mg/L]) compared to isolates without truncation of MmpR5, MmpL5 or MmpS5 (median 0.06 mg/L [stdev 0.1 mg/L]) (**Figure 6**). MmpL5 truncation appeared to cause hypersensitivity (median 0.008 mg/L [stdev 0.004 mg/L]), although we had few samples to assess this effect (n=4).

**Figure 6:**
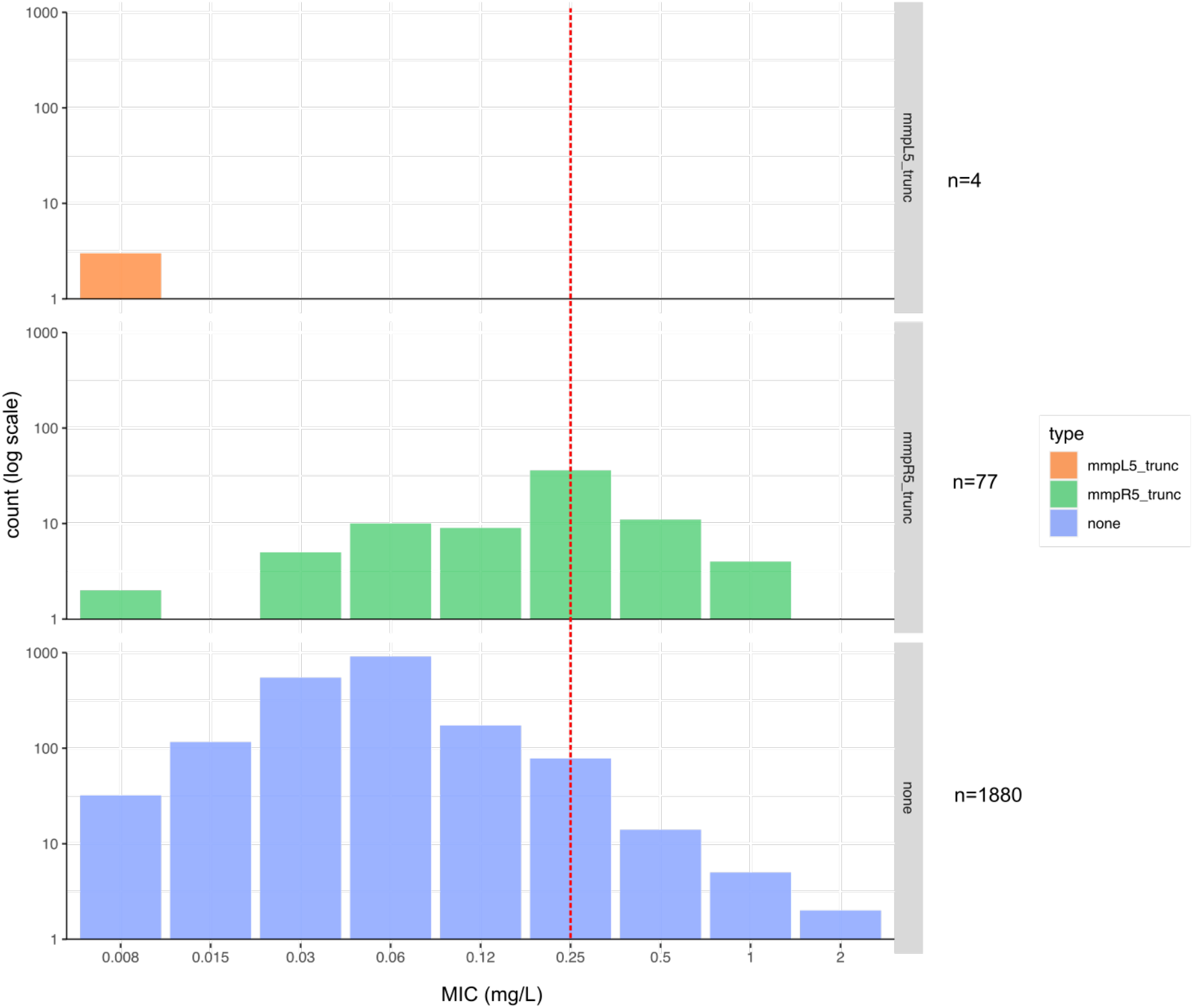
Bar plots of bedaquiline MIC based comparing truncation of MmpR5 and MmpL5 to no truncations. X-axis = bedaquiline MIC (mg/L), y-axis: count (log scale). Red dotted line: ECOFF cutoff (0.25 mg/L). mmpL5_trunc = isolates with MmpL5 truncated. mmpR5_trunc = isolates with MmpR5 truncated. None = isolates with no truncation of MmpR5, MmpL5 or MmpS5.

### *mmpR5* mutations led to a median 2-fold increase in BDQ MIC

We next tried to assess the contribution of each mutation in *mmpR5* to BDQ resistance by comparing the MICs of closely related pairs (at a threshold of 20 SNPs) where one contained a mutation in *mmpR5* and the other contained no mutations in *mmpR5, mmpL5* or *mmpS5*. Of 127 samples with mutations in *mmpR5* (including non-synonymous mutations, IS, nonsense mutations, and frameshifts), 58 samples were not assessed as they either had no closely related sample with MIC data within 20 SNPs (n=32) or the sample itself did not have MIC data (n=26).

Overall there was a median 2-fold increase in BDQ MIC in samples with mutations in *mmpR5* compared to closely related samples without mutations (**Figure 7**). The strongest effect overall was a frameshift at G66 (G66fs:SAPMP) which resulted in a 4-5-fold increase in BDQ MIC, although it was only found in two samples. The two positions with the highest number of samples (C46fs:GSRAA and P48fs:RAAVL) had variable results; the frameshift at C46 resulted in a median 2-fold increase in BDQ-MIC (ranging from 1-6 fold increase), while the frameshift at P48 also had a median 2-fold increase but contained 4 pairs (22%) with no MIC increase, and one pair with a 1-fold reduction in BDQ MIC. 18 out of 29 variant positions (62%) only had 1 pair that we were able to assess (**Figure 7**).

**Figure 7:**
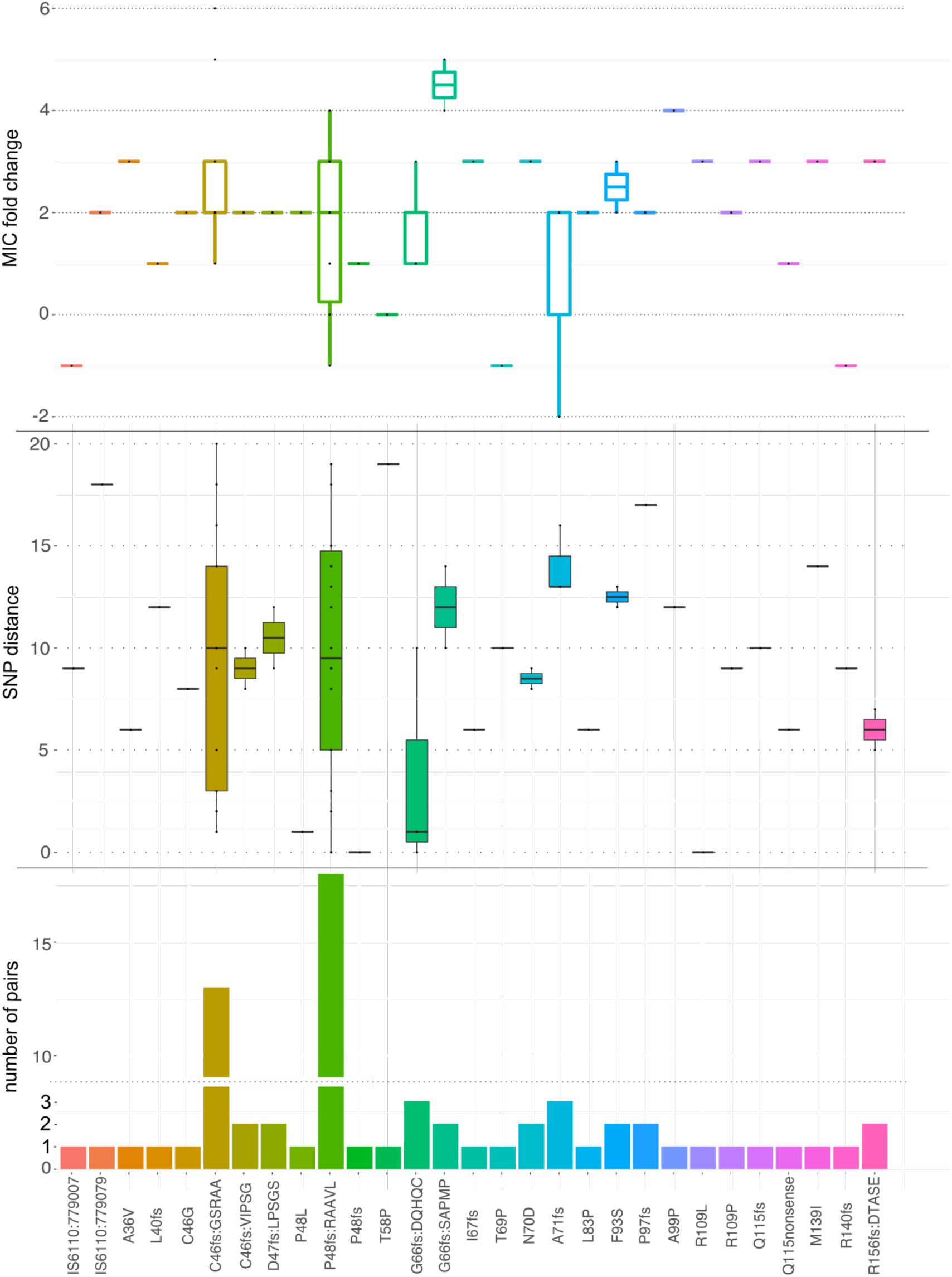
Estimated impact on MIC of observed variants in *mmpR5*. For each “case” isolate containing a variant (x axis) a control isolate is selected which does not have that variant, and which differs by at most 20 SNPs from the original isolate. The difference in MIC is a point estimate of the effect size. The distribution of effect on MIC across all isolates is shown in the top panel, the SNP differences between case and controls is in the middle panel and the number of case/control pairs is in the bottom panel.

## Discussion

BDQ resistance was first observed in 2005 and has since been seen repeatedly in South Africa, China, India, and elsewhere^2,13^. There are multiple mechanisms causing resistance; mutations in the target ATP synthase *atpE* cause extremely high levels of resistance but are clinically extremely rare^10^. It is more common to see mutations in the efflux pump regulator *mmpR5*, but so far it has not been possible to predict resistance with any great success; most mutations are unique and not consistently associated with resistance. In this study we analyse a collection of recent (2019-2020) samples from Cape Town and Johannesburg and find a notable enrichment of *mmpR5* truncations in BDQ-R samples. We then extend to a larger dataset consisting of 1,961 South African isolates with MICs, augmented with all other South African samples from the ENA to provide context.

In line with previous studies, we identified mainly lineage 2 and lineage 4 isolates, with few samples from lineages 1 and 3^16^. Using a strict threshold of 5 SNPs^17^ we found little evidence of recent person-to-person transmission in our cohort. There was also limited geographical clustering, with lineages identified across both sites (Johannesburg and Cape Town). The exception was a large number of lineage 2.2 isolates collected mainly from Cape Town, with a median 6 SNPs to the nearest neighbour (range 0-29 SNPs) indicating probable ongoing transmission at this site. We identified truncation of *mmpR5* as a strong indicator of increased BDQ MIC in our cohort and conclude that resistance evolved repeatedly across all lineages containing isolates with phenotypic BDQ-R and not as a result of transmission^18^. *mmpR5* has been the focal point of multiple BDQ-R-association studies^4,10^, but few have looked at truncation as an overall effect relating to BDQ-R. Our study also highlights the effect of *mmpR5* truncation in a sizable cohort of *M. tuberculosis* samples, including not only our primary SA1 dataset, but also a large number of publicly available *M. tuberculosis* samples with paired MIC data (SA2)^13^.

We assessed disruption of *mmpR5* using local assembly to simultaneously identify multiple means of truncation, including frameshifts, nonsense mutations, and also IS elements, which have been reported previously in separate studies^10,19^. We show that the most common mutations occur prior to the predicted DNA binding region of *mmpR5*, suggesting inactivation of the protein, although this remains to be confirmed *in vitro*.

We found very few cases of *mmpL5* or *mmpS5* disruption, and no evidence of combined inactivation of both the regulator *mmpR5* and the efflux pump *mmpL5*/*S5*. This suggests that inactivation of the efflux pump is rare (although not impossible^20^) and may convey a negative fitness cost to the bacterium. As such, detecting variation in *mmpR5* alone is not sufficient to predict BDQ resistance, and further work is required to understand expression levels of *mmpR5*/*L5/S5* and fitness costs associated with expression of this efflux pump *in vivo*.

We show that a majority of isolates with a truncated (or otherwise frameshifted) *mmpR5* had a median 2-fold increase in MIC to BDQ. This modest increase in MIC is in line with other studies looking at mutations in *mmpR5*^10^, and often places the BDQ MIC at the threshold of resistance (based on the ECOFF cutoff of 0.25 mg/L). This explains why binary phenotyping alone reports a mix of both sensitivity and resistance^21^.

While we found that truncations in *mmpR5* were important, we were limited in our ability to determine the specific contribution of certain mutations to BDQ susceptibility. Like other studies, we found a number of mutations, many of which were only represented by a single isolate^4,10^. For each isolate with a truncation of mmpR5 we used a phylogenetically closely related control without the truncation, to estimate the effect of the truncation on MIC. Most such pairs differed also by other SNPs than just those in *mmpR5*, making it difficult to assess the contribution of the *mmpR5* mutation alone.

In practical terms, we found that ARIBA is a robust tool for screening large datasets and identifying gene truncations, which was only slightly less sensitive than exhaustive manual curation. However, we found it highly sensitive to data quality (including at regions of high GC content^21^ especially in *mmpL5*), so recommend that predictions be separately and rigorously assessed.

We acknowledge some other limitations of this study. Firstly, we focused mainly on truncations and disruptions of the predicted protein sequences, and so did not rigorously assess synonymous mutations. While we looked for promoter disruptions in the SA1 dataset, we did not look for promoter disruptions in the SA2 dataset as we used ARIBA to screen genes only. Finally, we focused our IS analysis on IS6110 only, and did not explore other IS movement.

Overall, we recommend the continued use of MICs to identify modest shifts in BDQ resistance and local assembly tools, like ARIBA, to detect gene inactivations caused by truncations or disruptions. The impact of elevated MIC on patient outcome is also of great interest and warrants future study.

## Methods

### Study setting and sample selection

Isolates, cultured using the MGIT platform, included in this study were prospectively collected between 2019 - 2020 from two high-volume laboratories (Gauteng and Western Cape) in South Africa as part of surveillance activities in the Metropolitan areas. These were a minimum of MDR-TB by WHO definition.

### Culture, DNA extraction and whole genome sequencing

*M. tuberculosis* samples were cultured as previously described^13^. A 1.2ml aliquot of the cultured isolate was heat-inactivated at 80°C for 20 minutes in a forced air oven. The samples were pelleted by centrifugation at 8000 xg for 5 minutes, 600µL of the supernatant was discard and the resultant pellet homogenized by vortexing at maximum speed for 10 minutes in the remaining solution. DNA extraction was performed using 500µL of the sample on the NucliSENS easyMAG automated extraction system (Biomerieux, France). The Generic protocol was utilised with a final elution volume of 50 µL. Paired-end libraries were prepared using the Nextera Flex DNA library preparation kit (Illumina, USA) and sequenced using the either the MiSeq and NextSeq 550 instruments (Illumina, USA).

### Drug resistance phenotyping

Two-fold serial dilutions of 100x working solutions of BDQ (obtained through the NIH HIV Reagent Program, Division of AIDS, NIAID, NIH: Bedaquiline Fumarate, ARP-12702, contributed by Janssen Pharmaceuticals). A BDQ stock solution of 840 µg/ml was prepared by adding 10.08 mg BDQ powder (fumarate salts) to 10 ml dimethylsulfoxide (DMSO). These were stored at -70°C and on the day of testing, thawed at ambient temperature. A working solution was prepared by a 1:10 dilution with DMSO. DST was performed on the BACTEC™ MGIT™ 960 DST, a 100µl of the working solution was added to the MGIT tubes giving a final concentration of 1µg/mL (WHO tentative recommended critical concentration) and processed further as described previously* with minor modification, i.e. the incubation time was extended to 28 days.

### Quality control

All raw reads were run through the quality control and decontamination pipelines in Clockwork v0.10.0 (https://github.com/iqbal-lab-org/clockwork) using the reference genome H37Rv (accession: NC_000962.3). Samples that had <25x average coverage and/or >5% contamination were removed, leaving 505 *M. tuberculosis* samples (referred to as SA1).

### Variant calling and regenotyping

Per-sample variant calling was completed using Clockwork v0.10.0 (https://github.com/iqbal-lab-org/clockwork) against the reference genome H37Rv (accession: NC_000962.3). Clockwork is a variant caller which runs two other independent callers (SAMtools^22^, Cortex^23^) on each isolate, and then builds a graph genome of the (reference genome and) two sets of variant calls, and remaps the reads to this graph, in order to adjudicate between the two callers where they disagree^24^. Having done this, Minos^24^ v0.12.0 was used to collect a non-redundant list of all segregating variants in the cohort, harmonise variant representation in positions where indels overlap SNPs (or other indels), and then remap the decontaminated reads one more time, to genotype all samples at all variants, providing consistent VCFs for all samples.

### Publicly available South African TB

Raw sequencing reads for all publicly available South African *M. tuberculosis* samples from the European Nucleotide Archive (ENA) until October 2020 and the CRyPTIC dataset^13^ were downloaded (n=5,802). All samples underwent quality control (described above), leaving 5,253 *M. tuberculosis* samples (referred to as SA2). Specific isolate sets used for each analysis is given in **Supplementary Dataset 1**.

### In silico lineage assignment and resistance profiling

Mykrobe^25^ v0.9.0 was run on default settings using the raw reads against the 2020-01-14 TB panel. The output consisted of json files containing resistance predictions, mutation evidence, and lineage predictions.

### Phylogenetic reconstruction

Single nucleotide polymorphisms (SNPs) were substituted into the H37Rv genome using BCFtools^26^ consensus (v1.10.2) to create a whole-genome alignment of all samples (https://github.com/LeahRoberts/Mtb_South_Africa/tree/main/scripts). Repeat regions (*e*.*g*. PE/PPE genes) and genes known to be involved in antimicrobial resistance (including 100 base pairs upstream and downstream) were masked (https://github.com/LeahRoberts/Mtb_South_Africa/tree/main/mask_files) in the alignment. SNP-sites^27^ (v2.5.1) using the ‘-c’ flag was used to obtain all variant positions, as well as all constant sites using the ‘-C’ flag. The final alignment of variant position and constant sites were then used to generate a phylogeny with IQTree^28–30^ (v2.2.0 or version 1.6.12, see figure legend) and ultrafast bootstrapping using the parameters ‘-st DNA -nt AUTO -bb 1000 -m MFP’. Four variant sites with multiple alleles (150321, 55533, 976889, 3741263) were removed from the large SA1+SA2 South African *M. tuberculosis* phylogeny.

### Variant association with phenotypic data

To evaluate the contribution of specific mutations to BDQ MIC, we used a custom python script to combine the SNP distance matrix, ARIBA variant information (methods below), and isolate MICs (https://github.com/LeahRoberts/Mtb_South_Africa/tree/main/scripts).

### Investigation of bedaquiline-associated mutations

Several genes were investigated for their role in bedaquiline resistance based on previous literature, which we refer to as “bedaquiline-R-associated” genes (*mmpL5, mmpL3, mmpS5, atpE, amiA2, pepQ, era, rv1816, rv3249c, rv0678/mmpR5*)^2,13,31,32^. For the SA1 dataset, we performed *de novo* assembly of all isolates using Shovill v1.1.0 (https://github.com/tseemann/shovill) (which uses SPAdes^33^) with ‘--gsize 4.4M’ to investigate “bedaquiline-R-associated” genes variants and their effect on the predicted proteins. We then used blastn^34^ v2.9.0 and a custom script (https://github.com/LeahRoberts/Mtb_South_Africa/tree/main/scripts) to determine nucleotide similarity and the predicted amino acid translation of the “bedaquiline-R-associated” genes from the *de novo* assemblies. The predicted protein sequences were then aligned with MAFFT^35^ v7 at default settings and visualised using a neighbour joining tree created through Jalview^36^ v2.11.2.5. We used ISmapper^14^ v2.0 to characterise IS6110 insertion sites in the isolate genomes using the decontaminated reads and the reference genome H37Rv (accession: NC_000962.3).

To check for duplicated genes in our SA1 dataset, we mapped the decontaminated isolate reads to the H37Rv genome using BWA-MEM^37^ v0.7.17-r1188 and checked coverage across the entire genome and at specific gene sites (*mmpR5, mmpL5, mmpS5*) using pysamstats v1.1.2 (https://github.com/alimanfoo/pysamstats) and a custom script (https://github.com/LeahRoberts/Mtb_South_Africa/tree/main/scripts) to revise our conclusions to the appropriate rationale.

For the SA2 dataset, we used ARIBA^15^ v2.14.6 to broadly identify truncated or interrupted genes by creating a database consisting of a multi-fasta of the “bedaquiline-R-associated” genes, which were then used as the query database when running ARIBA. We then used ARIBA summary to concatenate all results and identify isolates where the gene prediction from ARIBA was “no”. Isolates with predicted truncations were analysed with Shovill, blastn, MAFFT, Jalview and ISmapper (as above). To control for poor quality data that could affect ARIBA’s local assembly, we did not consider predicted truncations where the sample’s Shovill assemblies generated >500 contigs. We also checked sample read quality using fastp^38^ v0.22.0 with the following parameters: “--detect_adapter_for_pe --dedup --length_required 50”. Samples with >20% of reads removed were not considered for predicted truncation events. Finally, we required predicted truncation events to be in agreement between the *de novo* assembly and the ARIBA result for that isolate.

A small number of isolates (n=4) had a truncated *mmpR5* but we were unable to determine the biological cause (no frameshift, no nonsense mutation, no IS, etc). This could be due to poor quality sequencing data (resulting in a fragmented assembly) or a biological event unresolvable with short-read sequencing (such as a large inversion, or repetitive elements). As we could not discriminate between these two events, we did not include results for these isolates in relation to BDQ-R.

### Plotting and figures

We used iTol^39^ to visualise all phylogenies. We used ggplot2^40^ with the plyr^41^ package to visualise the BDQ MIC distributions.

## Contributions

LWR, ZI, SVO and MF contributed to the conceptualisation of the manuscript. SVO and LJ contributed to the data collection and antimicrobial susceptibility testing. DC contributed sequencing of the samples. LWR, KMM, PW, MH and JK contributed to the sample processing. LWR, PW and ZI contributed to the methodology and formal bioinformatic analysis. LWR contributed to the writing (original draft). ZI, KMM, MH and MF contributed to the writing (review and editing). ZI, SVO and MF contributed to the supervision of the study. All authors had access to the data presented in this study and had final responsibility for the decision to submit for publication.

### Data sharing

Study samples and accessions are available in **supplementary dataset 1**.

### Funding

LWR was supported by an EMBL Biomedical Postdoctoral Fellowship (EBPOD). Financial support for next generation sequencing from the CDC – USA (NU2GGH002194)

## Acknowledgements

The authors would like to acknowledge the CRyPTIC consortium for their collection and open release of *M. tuberculosis* WGS samples with matched MIC phenotypes.

