## Supplementary Material for "Repeated evolution of bedaquiline resistance in *Mycobacterium tuberculosis* is driven by truncation of *mmpR5*"

### Supplementary results:

*Notable clonal expansions from SA1 dataset:*

The SA1 phylogeny revealed expansion of 183 isolates from lineage 2.2, primarily from Cape Town, with a median SNP distance of 6 to the closest isolate (average = 7.7 SNPs, range = 0-29 SNPs) (**Supplementary Figure 1**). We also identified 20 highly resistant isolates from lineage 2.2.1, with resistance to isoniazid, rifampicin, quinolones (ofloxacin, moxifloxacin, ciprofloxacin), aminoglycosides (kanamycin, amikacin, capreomycin, streptomycin), ethambutol, and pyrazinamide (**Supplementary Figure 1**). These resistant isolates were found in three phylogenetically distinct clusters from both Cape Town and Johannesburg, that on average differed by 18-21 SNPs within each cluster.

### Supplementary Figures:

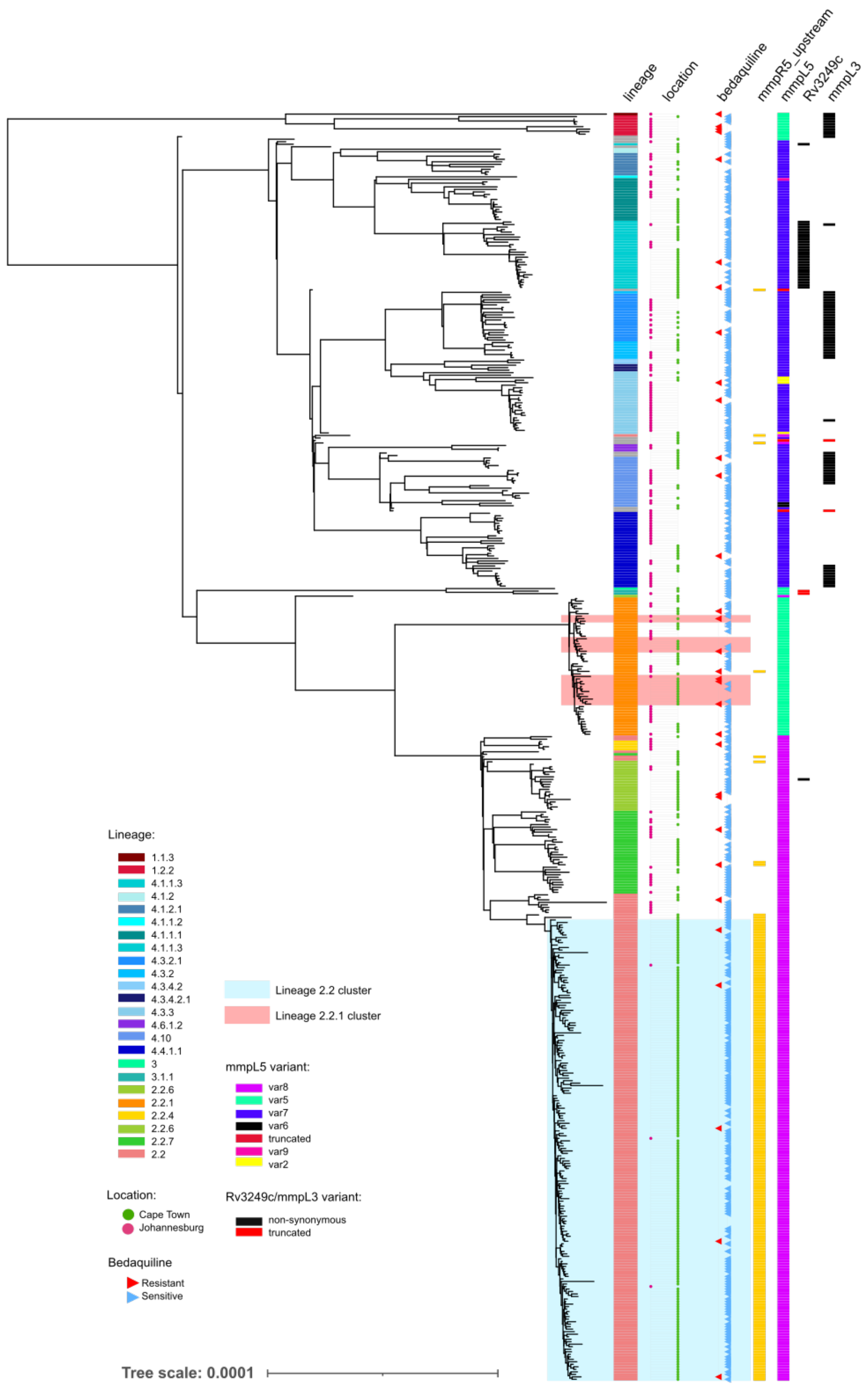


**Supplementary Figure 1: phylogenetic clusters and mutations:**
mmpR5_upstream: any mutation (compared to H37Rv) in the upstream region of mmpR5. mmpL5 variant: non-synonymous variant type (or truncation [red]).

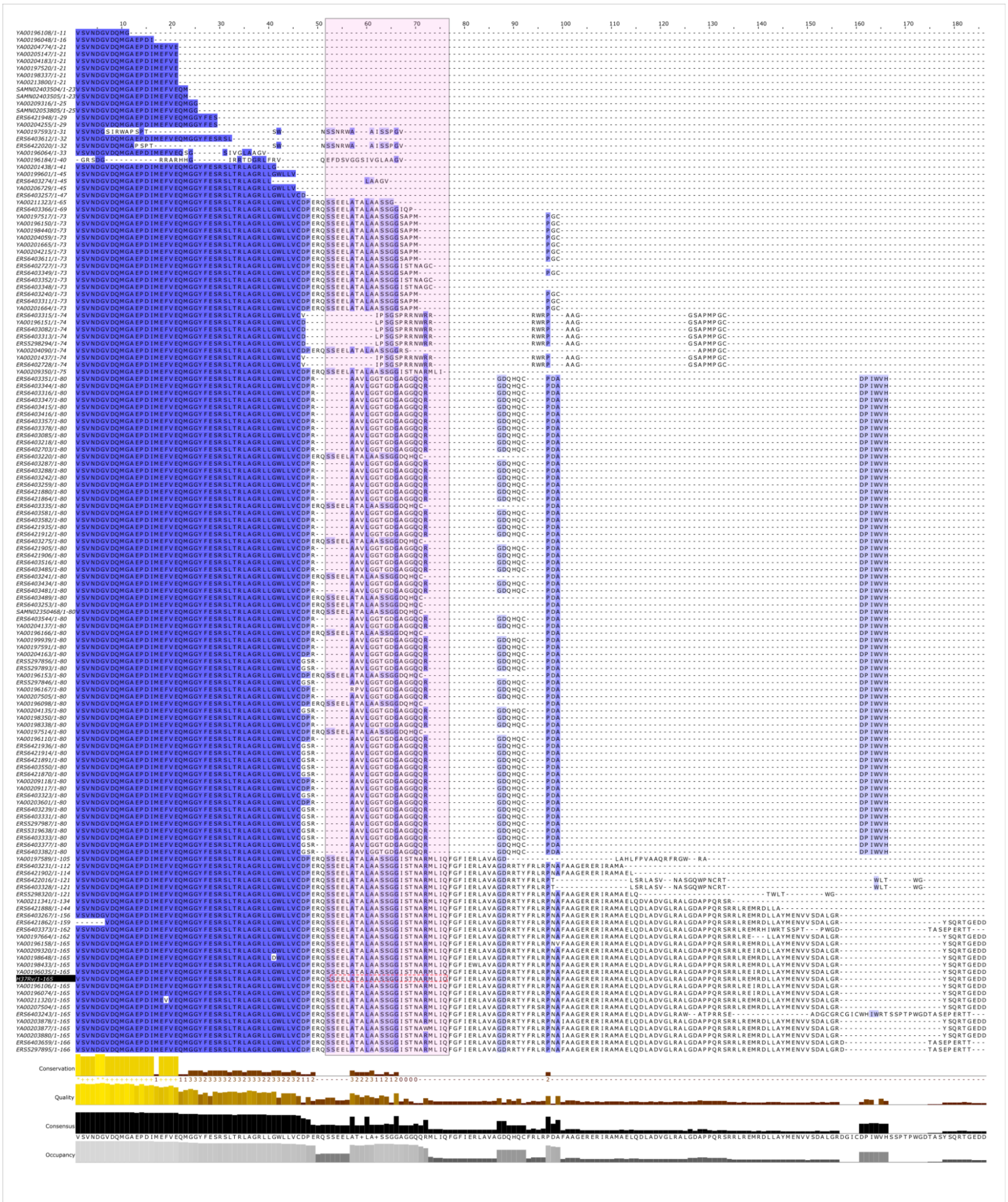

**Supplementary Figure 2**: MAFFT protein alignment of MmpR5 predicted sequences from all isolates with truncated MmpR5, and SA1 dataset isolates with non-synonymous amino acid substitutions. Purple (spectrum) represents amino acid similarity at that position of the alignment (darker = higher similarity). Light pink area is the DNA-binding domain for MmpR5. Lower barplots: Conservation: numerical index reflecting the conservation of the physico-chemical properties, Quality: ad-hoc measure of the likelihood of observing the mutations, Consensus: the percentage of the different residue per column, Occupancy: the number of residues aligned at each column
